# Sensory context of initiation-cue modulates action goal-relevant neural representations

**DOI:** 10.1101/2024.09.03.611077

**Authors:** Nicholas Kreter, Neil M. Dundon, Jolinda Smith, Michelle Marneweck

**Affiliations:** University of Oregon, Department of Human Physiology, Eugene, OR, 97403; University of California Santa Barbara, Department of Psychological and Brain Sciences, Santa Barbara, CA, 93106; University of Freiburg, Department of Child and Adolescent Psychiatry, Psychotherapy and Psychosomatics, Freiburg, 79104, Germany; Institute of Neuroscience, University of Oregon, Eugene, OR, 97403; Phil and Penny Knight Campus for Accelerating Scientific Impact, Eugene, OR, 97403

**Keywords:** Reaching, Neural Representations, Multisensory, fMRI

## Abstract

The ability to produce goal-directed movement relies on the integration of diverse sources of sensory information specific to the task goal. Neural representations of goal-relevant features, such as target location and gaze direction, have been well studied in sensorimotor areas. It remains less clear whether goal-relevant motor representations are influenced by sensory changes to initiation-relevant information, such as a go-cue that provides no information about target location. We used Bayesian pattern component modelling of fMRI data during a delayed reach task with either visual or audiovisual go-cues to explore whether neural representations of goal-related features in sensorimotor areas are modulated by changes to initiation-relevant sensory information. We found that representations of target direction and gaze direction in the primary sensory areas, motor areas, and posterior parietal cortex, were sensitive to whether a reach was cued with a visual or audiovisual go-cue. These findings indicate that the central nervous system flexibly delegates the tasks of ‘where’ to move and ‘when’ to move based on available sensory context, even if initiation-relevant stimuli provide no additional information about target location.

## 1. Introduction

The ability to perform goal-directed movements is essential to many behaviors in daily life. Successful execution of these actions often necessitates integrating rich sources of sensory stimuli in support of movement subtasks, such as tracking when to initiate a movement in the presence of an external cue^1^, and monitoring the position and orientation of the necessary effector relative to a target location^2,3^. For instance, the simple goal-directed action of reaching to a phone in response to an incoming call involves contributions from visual, proprioceptive, and auditory systems. The visual system helps to identify the phone’s location in space relative to the body, the proprioceptive system helps identify the position and orientation of the hand that will be used to reach, and if the phone is ringing, the auditory system will provide information about the spatial location of the phone and an external cue that the reach should occur.

During externally cued actions, sensory systems gather goal-relevant information (i.e., information about where or how to move) or initiation-relevant information (i.e., information about when to move, absent of spatial information). Both goal-relevant and initiation-relevant information can be relayed to the brain by various sensory systems. When an incoming call lights up the phone screen, visual stimuli travel from the primary visual cortex (V1) to the posterior parietal cortex (PPC) via the dorsal visual stream, providing both goal- and initiation-relevant reach information. If the screen lights up and a ringtone plays through a set of headphones, the visual system receives both goal- and initiation-relevant information while the auditory system receives only initiation-relevant information. In this instance, auditory stimuli traveling from the primary auditory cortex to PPC do not specifically provide information about where or how to reach, but its availability allows a division of labor for the goal-relevant and initiation-relevant subtasks when preparing an action. That is, the goal-relevant subtask of localizing a target (i.e., where and how to move) may be assigned to the visual system, while the auditory system could monitor the subtask of movement initiation (i.e. when to move).

It is well established that the sensory context supporting a goal-directed action changes neural representations in sensorimotor areas. Visual, auditory, superior parietal, dorsal premotor, and primary motor areas show distinct goal-relevant representations of reach targets that are presented with visual^2-5^, auditory^6-8^, or proprioceptive^9^ cues, suggesting that these regions are flexible to changing sensory contexts of goal-related sensory information. Despite the abundance of research exploring how goal-relevant reach information is flexibly represented within sensorimotor pathways, it is less clear how initiation-relevant information is represented within the same regions, and whether sensory changes to initiation cues modulate the representations of goal-relevant reach information.

Much of the research exploring the neural representations of reaching has focused on understanding how the brain represents the goal-relevant structure of reaching with less regard to how the brain represents initiation-relevant structure of reaching. The dorsal visual and dorsal auditory streams are often referenced as ‘where’ or ‘how’ pathways^10^, with less consideration for how these structure handle when to move^11^, or how these processes interact. However, given that the dorsal stream has a temporal advantage over the ventral stream ^12,13^, it is unsurprising that it would also handle temporal cues. Several studies have suggested that the dorsal auditory stream plays a key role not only in spatial location, but also in timing when events happen^11,14,15^.

Further, pre-motor and PPC areas are responsive to temporal uncertainty in simple and choice reaction time tasks^16^, suggesting that the same regions involved in monitoring where to move also monitor when to move.

Here, we sought to verify that goal-relevant and initiation-relevant information interact within sensorimotor regions by investigating how neural representations of visually guided reaching change when performed with different initiation cues. Participants performed a reach to target task across two sessions where they 1) had only visual information guiding movement initiation, and 2) audiovisual information to guide movement initiation. Given the evidence for sensorimotor brain regions being sensitive to spatial and temporal task structure, as well as the sensory context in which a task is performed, we hypothesized that changes to initiation-relevant sensory cues would modulate the goal-relevant representations of reaching.

## 2. Materials and Methods

### Participants

Twenty-seven healthy young adults [17 f, mean (SD) = 21.3 (3.2) years] were recruited from the local community and provided informed consent to participate in this IRB approved study. Exclusion criteria included self-reported left-handedness (or as determined by the Edinburgh handedness inventory), neurological diagnosis, neuromuscular injury, and self-reported motor or cognitive impairment that would adversely impact the ability to perform the experimental reaching tasks.

### Materials, Design, and Procedures

Subjects completed a goal-directed reaching task using a custom-built task board during two counterbalanced scanning sessions. Sessions were performed on separate days and each session featured six functional runs. Structural imaging was performed during each participant’s first session. The task board featured LED lights that indicated the starting hand position, gaze direction, and reach target (Figure 1). Eight unique conditions were used that featured a combination of left or right initial hand position, gaze direction, and reach target direction. Participants performed these conditions across two sessions where only the sensory modality of the go-cue differed: one session featured trials with only visual go cues while the other featured an audiovisual go cue. During trials with the visual-only go cue, the gaze light changed color, and the target light turned off simultaneously. The trials with audiovisual go cues featured an auditory beep at the same time as the target light turned off.

**Figure 1.**
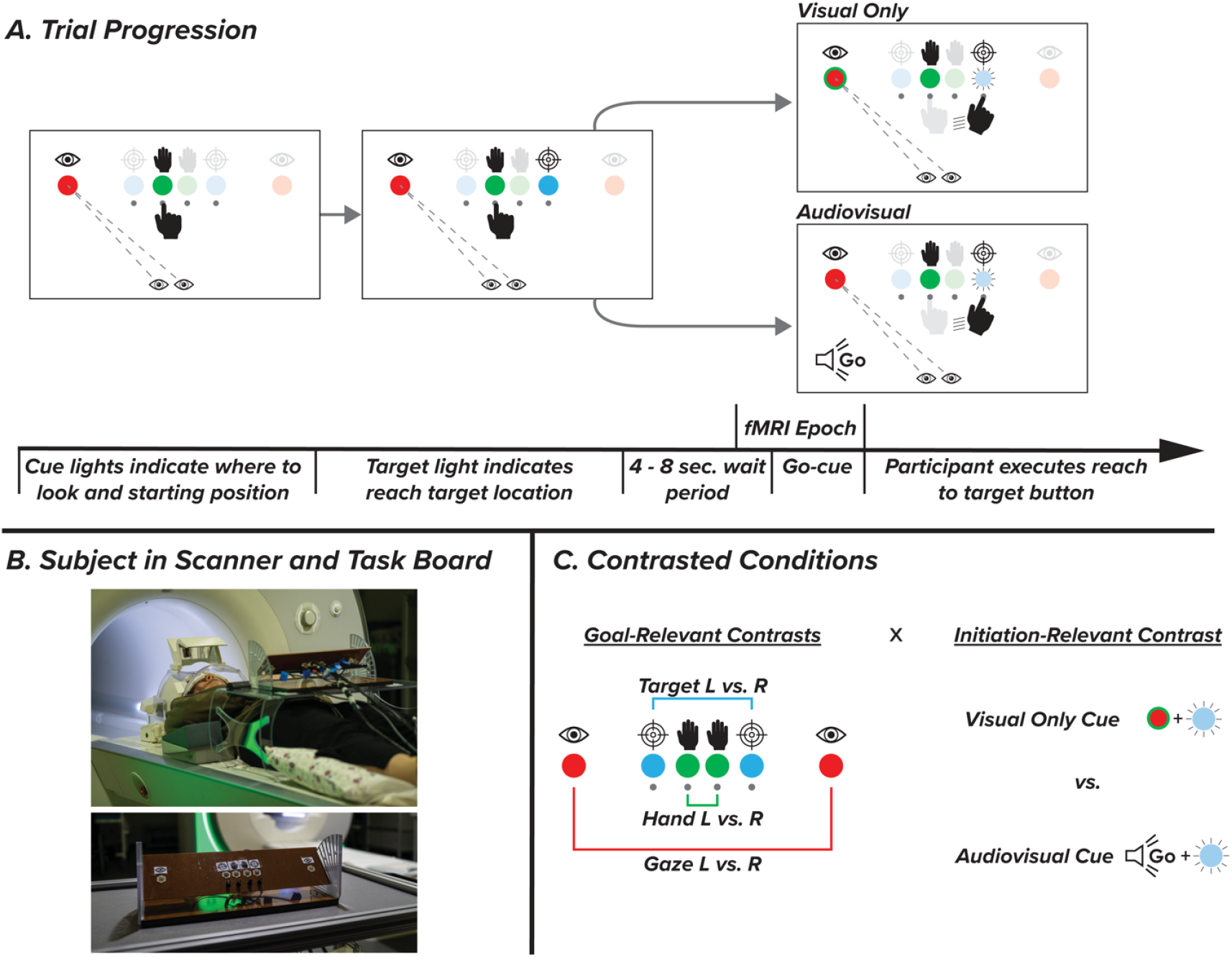
Experimental design and contrasts. A. Participants performed eight trial types with either left or right gaze, left or right initial hand position, and a left or right target. Trials started with a gaze light instructing participants where to look and a hand light instructing which button to start pressing with their right hand. Once pressing the correct button, a target light would appear. After a variable wait period of 4-8 seconds, participants received either a visual or audiovisual go cue and reached to the button under the target light. General linear models were computed to predict BOLD activity during the epoch covering the final two seconds of the wait period through movement initiation. B. Participants performed the task on an interactive board while laying in the scanner. A mirror attached to the head coil allowed participants to see each of the buttons, lights, and their hand. C. Contrasts tested the interaction between goal-relevant (gaze, target, and hand direction) and initiation-relevant (initiation-cue modality) task features, as well as individual main effects of direction and modality.

Each trial began when the LED indicating hand position and gaze direction illuminated. After participants pressed the correct hand-position button there was a variable delay period, followed by a variable plan period where a third LED indicated the reach target. The variable delay period was randomly drawn from a selection of [1, 2, 4, 8, or 16 seconds] with respective proportions of trials [0.52, 0.26, 0.13, 0.06, 0.03] while the plan period was randomly drawn from [ 4, 6 or 8 seconds] with respective proportions of [0.56, 0.30, 0.14]. Following the plan period, participants received a cue to reach towards the target. At the onset of the movement initiation cue, the hand and target light disappeared synchronously with the change to the color of the gaze LED (visual session), or the auditory cue played through headphones (audiovisual session). Following completion of the reach to target button, an audio cue played through headphones indicating success or failure. If the wrong target button was pressed, or no movement was made for 2.5 seconds after the go cue, the trial was counted as a failure. Each session featured six functional runs of 40 trials, with breaks between runs. In this study, a subset of trials featured a no-go stimulus where participants were instructed not to perform a reach, but data from these no-go trials are not presented in this manuscript. All events from stimulus presentation to button lift and button press timing were controlled with a custom Python script.

The order of trials and durations of the delay and plan periods were determined to minimize the variance inflation factor (VIF) within each functional run^17^. VIF estimates the degree of multicollinearity in a model by providing an index of how much the variance of a single regression coefficient increases due to collinearity. VIF values close to 1.0 indicate no change in variance between regressors whereas larger VIF values indicate that regressors are not independent of each other. In the present study average VIF was around 1.15, which indicates the independence of regressors^17^.

The dimensions of the task board were 40.0 cm wide, 10.0 cm long, and 6.5 cm deep. It consisted of the six LED lights indicating gaze, hand, and reach target locations, as well as four buttons beneath the hand and target LEDs. Icons indicating the gaze, hand, and target LEDs were placed above each LED. The LEDs indicating left and right-hand position were located 2.3 cm apart at the midline of the board. Each target and gaze LED was 2.3 cm and 6.9 cm lateral to each hand LED, respectively. Throughout testing, the task board was positioned on a stand 4 cm above the surface of the table and angled at 40.6 ° relative to the table surface. Pilot testing indicated that this position and angle were optimal for participants to maximize their view of the task. After receiving task instructions, but before being placed in the scanner, each participant performed practice trials prior to familiarize themselves with the experimental task.

All trials in this experiment were performed during fMRI in a Siemens Skyra (Siemens Medical, Germany; 32-channel phased-array head coil). During the first session, imaging began with a high-resolution T1 weighted anatomical scan (TR/TE = 2500/2.98 ms, flip Angle = 7°, FOV = 256 mm). Following the anatomical scan, participants performed the reaching task while BOLD contrast was measured with a multiband T2*-weighted echoplanar gradient echo imaging sequence (TR/TE = 450/30 ms, flip Angle = 45°, FOV = 192 mm; multiband factor: 6). Each full brain functional image consisted of 48 slices acquired parallel to the AC-PC plane (3 mm thickness; 3 × 3 mm in-plane resolution)^18,19^. To minimize the effects of motion, padding was added to secure the head, neck, and shoulder for all participants. The goal-directed reaching task was performed on an interactive board placed at arm’s length on a marked spot on a table spanning each participants’ hips. With a mirror attached to the head coil, participants could see the task board and their hand as if they were viewing them while sitting upright.

### Data Processing and Statistical Analysis

MRI pre-processing and analysis were performed with SPM12 (Welcome Trust Center for Neuroimaging, London, UK), FSL^20^, and custom MATLAB (Natick, MA) code. Using 2^nd^ degree B-spline interpolation in SPM, subjects’ functional images were spatially realigned to a mean image. Images were then coregistered to the T1-weighted image and normalized. Mean head motion rotations and translations were minimal (minimum and maximum values in parentheses): x: 0.015 mm (−2.681, 1.714); y: 0.108 mm (−1.886, 3.754); z: 0.215 mm (−3.922, 4.188); pitch: -0.0002° (−0.1471, 0.1509); roll: 0.0008° (−0.0365, 0.0387); yaw: 0.0013° (−0.0941, 0.0495). Functional images were each inspected for distortions and inhomogeneities.

Following pre-processing of the anatomical and functional imaging data, we assessed spatial patterned activity in pre-determined sensorimotor regions of interest (ROIs), including primary visual (V1) and auditory (Heschl) areas, superior parietal areas 5 (SPL5) and 7 (SPL7), PMd, and primary motor area 4a from the Julich brain atlas^21^. Spatial patterned activity differences were assessed between conditions in the ROIs by implementing Bayesian representational similarity analyses (vRSA)^22^. Convolution-based general linear models were run for each of the eight experimental conditions from each session. The general linear model specifically focused on event-based activity in a time window starting two seconds prior to the go-cue and ending at movement onset for each trial that participants performed successfully. We also modelled task error and activity related to movement, such as when participants reached to the specified target or when they returned their finger to the starting position.

Variational representational similarity analysis (vRSA) compared between-condition dissimilarity in spatial activity patterns in sensorimotor ROIs through a method that decomposes second-order statistics. The analysis procedure starts with a sixteen-row condition-by-voxel matrix (U) for each participant. These sixteen rows cover two initial hand positions (left vs. right), two gaze directions (left vs. right), two target directions (left vs. right) and two cue modalities (visual vs. audiovisual). This U-matrix accounts for the mean voxel activity pattern during each condition, as calculated by the general linear model. Next, we construct a sixteen-by-sixteen second-order similarity matrix (G = UU^T^) where the main diagonal represents the covariance explained by each condition. In this G-matrix, greater off-diagonal values indicate greater pattern similarity between conditions. Following the construction of the G-matrix, we can then test specific hypotheses by examining the contribution of components (i.e., effects) to G. Components included the initial hand position (left vs. right), gaze direction (left vs. right), target direction (left vs. right), initiation cue modality (visual vs. audiovisual), and the two-way interactions between hand, gaze, target location, and cue modality, respectively. Three-way interactions were included in the analyses but are not presented as they produced no evidence of a meaningful effect. Four-way interactions were not included. Visual representation of the ROIs and specific contrasts can be found in Figure 2.

**Figure 2.**
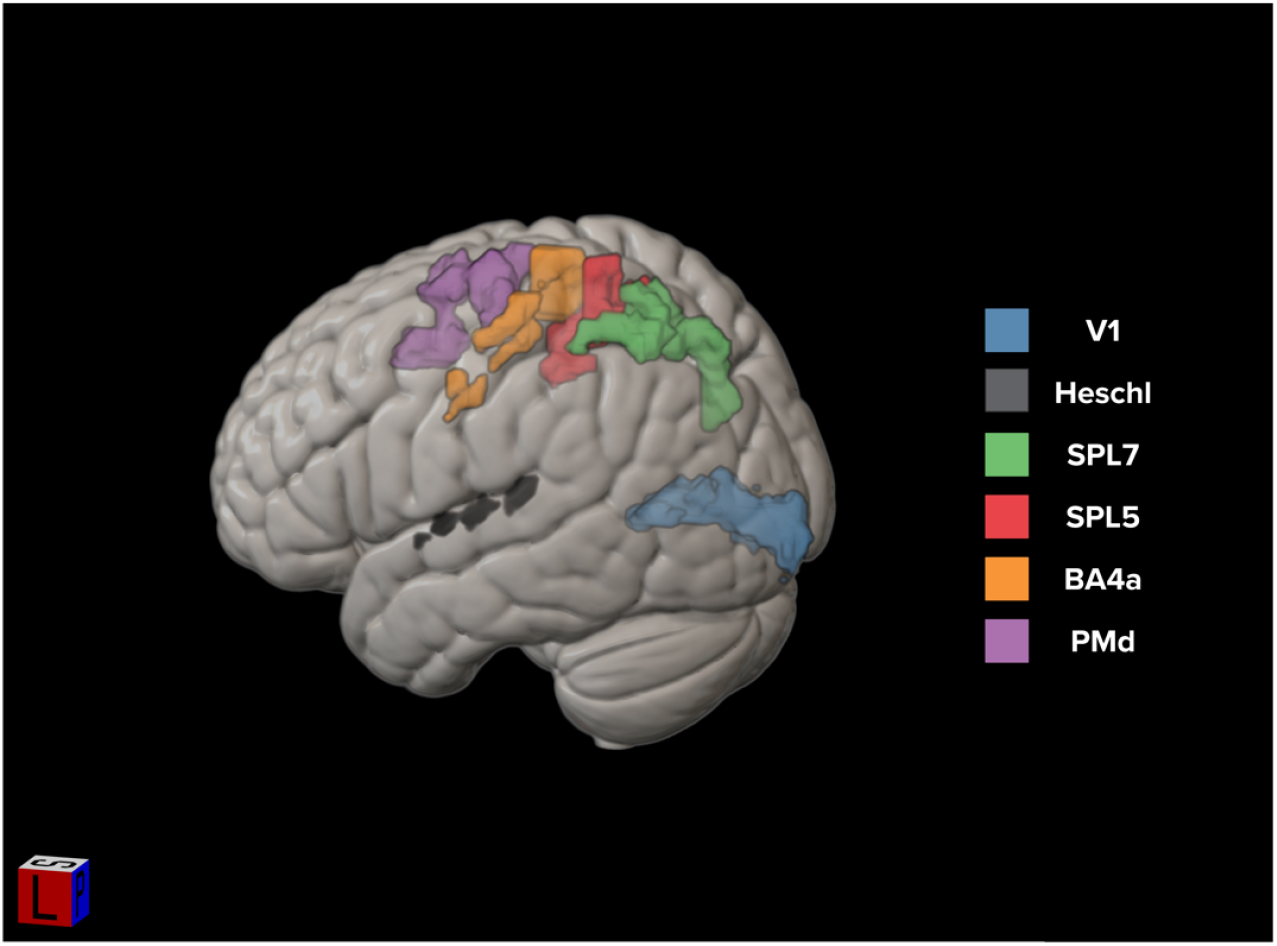
Predefined regions of interest from the Julich Brain Atlas are displayed on the MNI-152 atlas using MRIcroGL software. V1: primary visual, Heschl: Heschl’s gyrus, SPL: superior parietal lobule, PMd: dorsal premotor, BA4a: primary motor.

A strength of vRSA is assessing the independent contribution of each component to the covariance matrix G while controlling for each of the other contrasted conditions which the ROI may also be sensitive too. For example, if we observe an effect of cue modality in a specific ROI, that finding is true irrespective of whether the participant was planning to reach to a left or right target. Further, interaction effects in this analysis show whether changes in spatial patterned activity from one effect are dependent on changes to a second effect, irrespective of changes in each of the other effects. For example, a credible cue modality by gaze-direction interaction indicates that spatial patterned activity differences between left and right-sided gaze conditions vary depending on the sensory modality of the initiation cue. In other words, gaze direction representations are modulated by audiovisual and visual initiation cues (regardless of whether the hand or target was on the left or right). Any interaction between goal-relevant and initiation-relevant effects would confirm our hypothesis (i.e., sensory initiation cue * target direction, sensory initiation cue * gaze direction, or sensory initiation cue * initial hand position). These interactions would support the hypothesis that changes to initiation-relevant sensory cues modulate the goal-relevant representations of reaching. While the individual effects testing target direction, gaze direction, hand position, and cue modality do not directly test our hypothesis, we include them in our model because previous literature suggests existent representations in the tested sensorimotor ROIs. A specific benefit of the vRSA method is the ability to test individual hypotheses related to the contribution of a single effect, or interaction effects, while controlling for all other component effects included in the model comparison framework.

vRSA analyses performed using SPM return log evidence values of a hypothesis-specific representation at the group level. Log evidence values that are more negative indicate greater evidence of a credible effect. To ensure sufficient log evidence is observed, we performed an additional permutation analysis with 1000 iterations where condition labels were shuffled randomly. Calculating a G-matrix for each iteration and testing the shuffled data for the same main effects and interactions results in a null distribution for each contrast. We then subtract the actual (unshuffled) log evidence from the log evidence of each of the observations in the null (shuffled) distribution to create a distribution of log Bayes factors for each contrast, where higher values now communicate strong effects. We report whether the 95% highest density interval (HDI; analogous to a confidence interval) of the distribution of log Bayes factors is greater than log(3). Effects that meet this criteria are taken as being strongly credible results. In other words, we assume substantial evidence if the real data is three times more likely than the top 5% of data from the null distribution.

Our statistical methods adopt several distinct features to account for issues related to multiple comparisons. First, we use a hierarchical model with the same set of default priors to test each of the contrasted effects in parallel. Therefore, each model is an independent test of the pattern composition of an ROI. Second, for each ROI we performed a permutation analysis to create a null distribution of evidence specific to each contrast. While typical Bayesian analyses compare H1 relative to H0, this permutation analysis allows us to compare the H1 from the actual data, to the H1 of 1000 permutations of null data. We only conclude there is substantial evidence for condition-specific activity within a region if the H1 from the actual data is greater than the top 5% of data from the null-distribution. This step is a conservative addition, not typically included in vRSA analyses and diminishes the need for additional corrections due to multiple comparisons. See Marneweck et al. 2023^23^ for a comprehensive discussion on how this conservative approach mitigates the need for multiple comparison corrections within and across ROIs (also see ^24-27^).

## 3. Results

We used Bayesian pattern component modeling of fMRI data to assess whether goal-relevant representations in sensorimotor ROIs were sensitive to changes in initiation-relevant sensory cues during a goal-directed reaching task. Results largely supported our hypotheses that the sensory context of initiation-relevant cues modulates goal-relevant representations in sensorimotor ROIs. Specifically, we found broad evidence for goal-relevant representations of reach target location and gaze direction to vary depending on whether movement initiation cues were visual or audiovisual. However, for the effect of hand position, we found no evidence of goal-relevant representations in sensory or motor areas and no interaction between the effects of hand position and cue modality. These changes to spatial patterned activity with different initiation cues were independent of behavioral differences between trials with visual initiation cues versus those with audiovisual initiation cues.

### Gaze, Target, and Hand Interactions with Initiation Cue Modality

All regions showed effects for initial gaze direction and initiation cue modality, indicating that spatial pattern activity is distinct for left versus right gaze directions when movement initiation is cued visually versus audiovisually, respectively. We found strongly credible gaze*cue modality interactions in all regions [P(log BF > log(3)) ≥ 0.99]. The interaction between gaze direction and initiation cue modality indicates that differences in the patterned spatial activity between left and right gaze directions vary based on the modality of the go-cue.

Similar to the gaze*cue modality interaction, a target*cue modality interaction indicates that the differences in spatial patterned activity for changes in target location (left versus right) are modulated with different go-cue modalities. We found strongly credible target*cue modality interaction in all regions [P(log BF > log(3)) ≥ 0.99] except for V1 [P(log BF > log(3)) = 0.92]. While the results in V1 do not meet our criteria for strong credibility, and should not be taken as strongly credible, it should be noted that V1 falls just short of the threshold for strong credibility. Unlike frequentist statistical methods, which use sharp cutoffs to assess significance, our analysis provides a continuous measure of probability relative to the null data. Overall, the observed interactions between initiation cue modality and gaze direction, and target direction, suggest that representations of goal-relevant factors in reaching tasks are dependent on initiation-relevant factors, even if those factors provide no additional information about the required movement (Figure 3).

**Figure 3.**
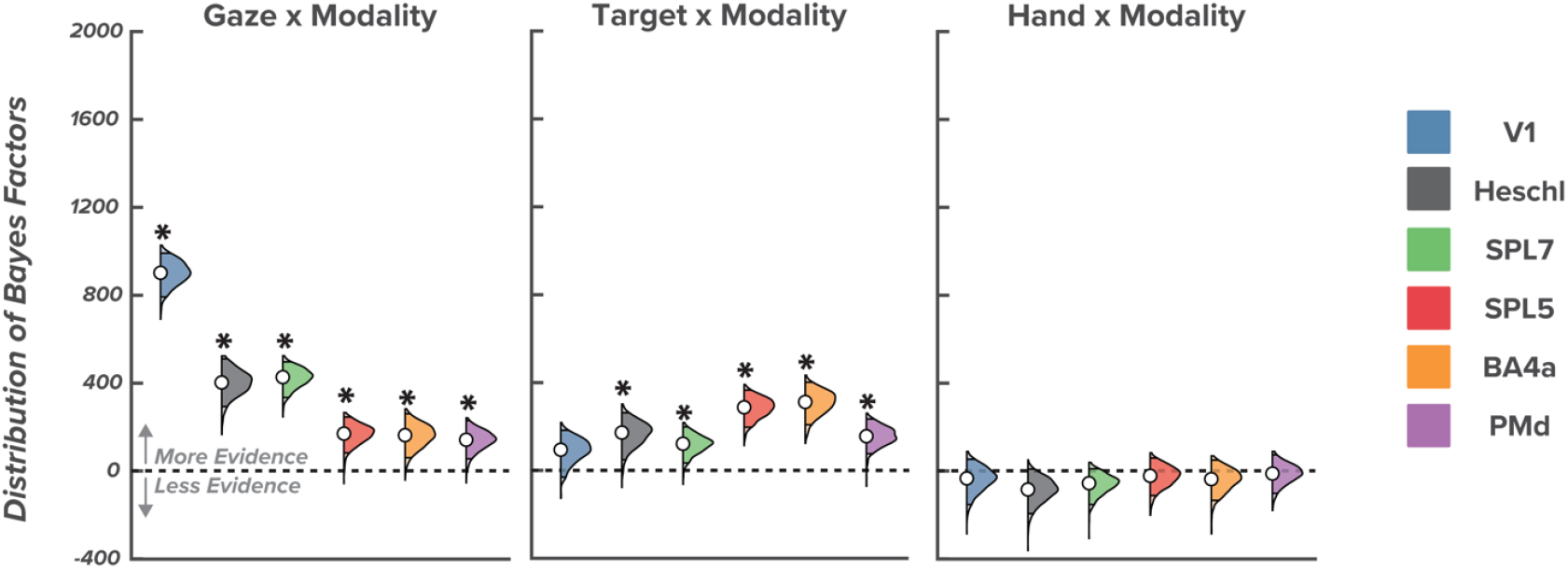
Interaction effects between goal-relevant and initiation-relevant effects. Violin plots show the distribution and 95% highest density intervals (HDI; dark-shaded region) of Bayes factors for the interactions between goal-relevant and initiation-relevant task features for each region of interest. Asterisks indicate substantial evidence that an effect is 3 times more credible than the 95% strongest effect from the null distribution in a specific region of interest.

**Figure 4.**
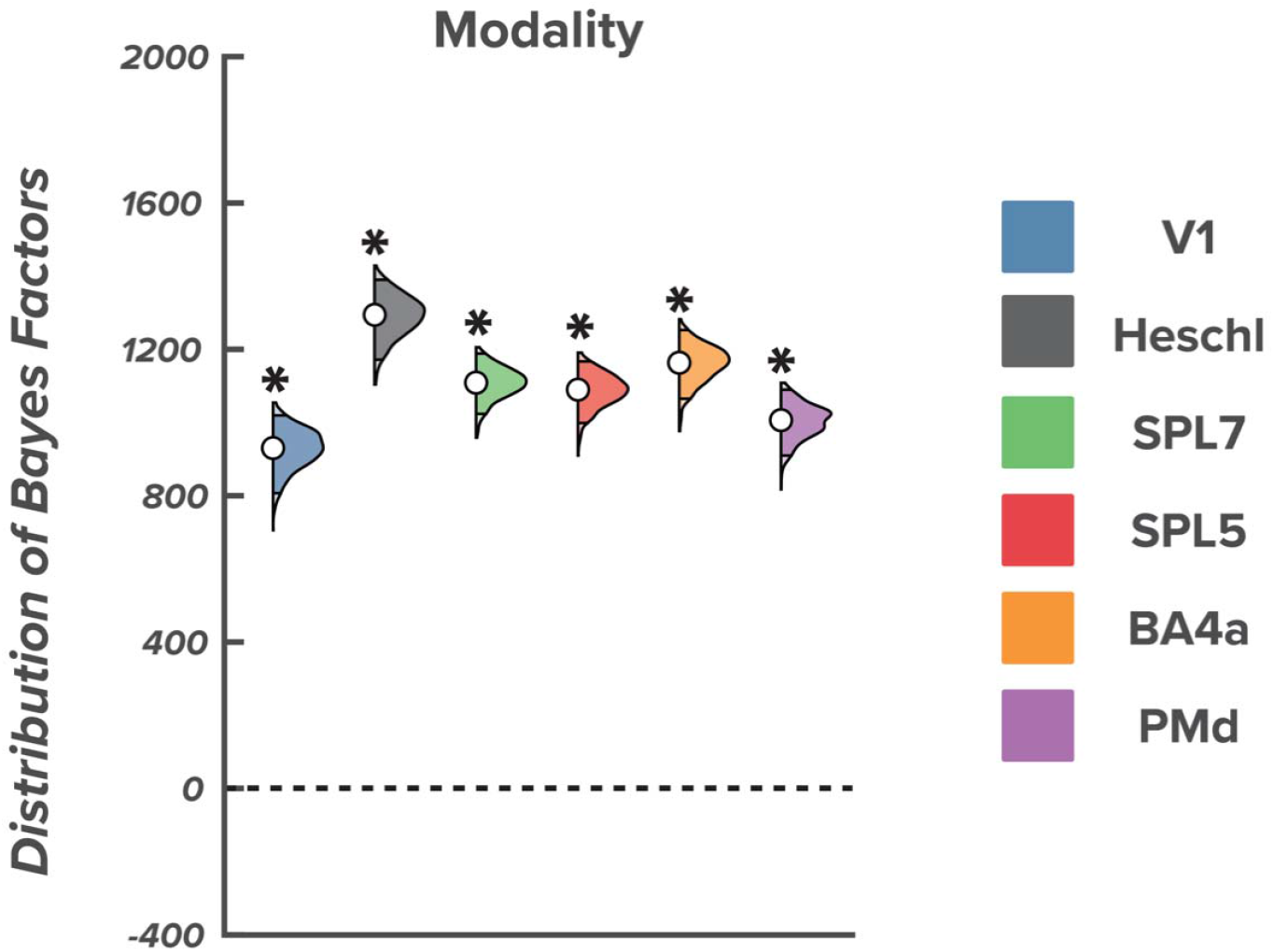
Effect of Initiation Cue Modality. Violin plots show the distribution and 95% highest density intervals (HDI; dark shaded region) of Bayes factors for the initiation-relevant effect of cue modality for each region of interest. Asterisks indicate substantial evidence that an effect is 3 times more credible than the 95% strongest effect from the null distribution in a specific region of interest.

**Figure 5.**
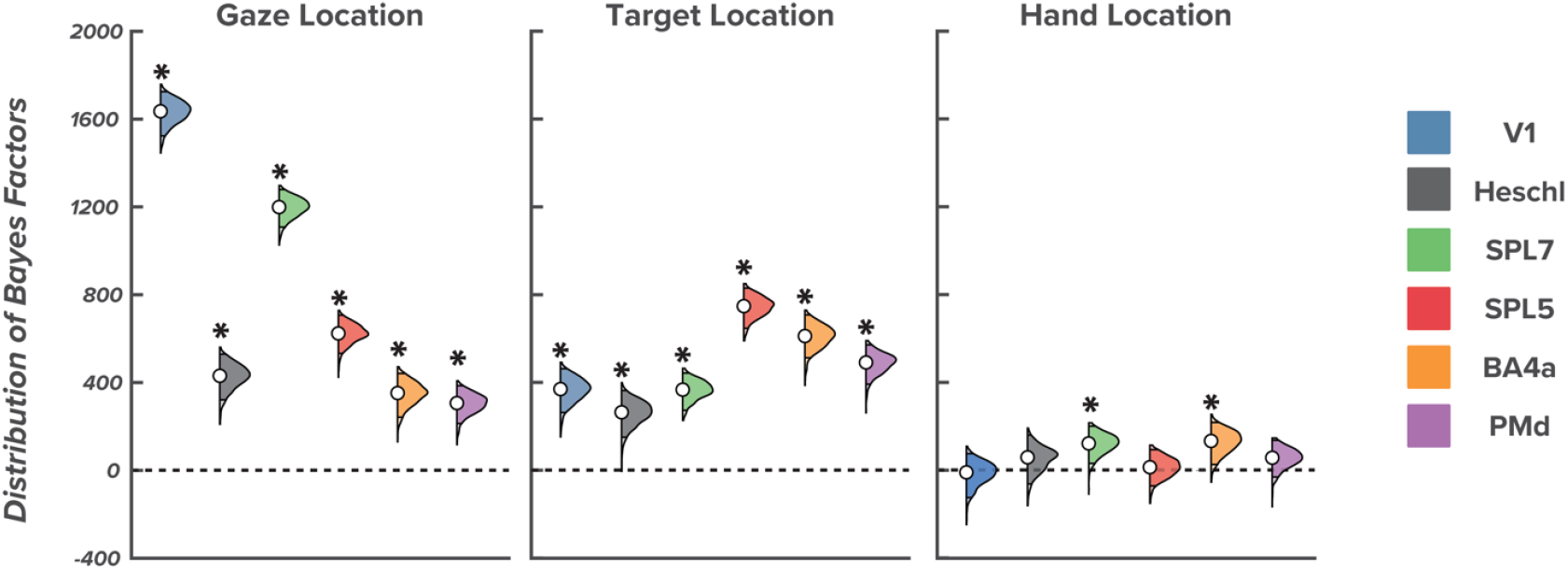
Effect of hand, gaze, and target location. Violin plots show the distribution and 95% highest density intervals (HDI; dark shaded region) of Bayes factors for the goal-relevant effects of gaze, target, and hand direction, for each region of interest. Asterisks indicate substantial evidence that an effect is 3 times more credible than the 95% strongest effect from the null distribution in a specific region of interest.

**Table 1.** Probability of credible evidence for each main effect and interaction in every ROI. ROIs in which P(BF>log(3)) exceeds 0.95 indicates that there is strongly credible evidence for distinct representations.

|  | Hand | Gaze | Target | Modality | Hand x<br>Modality | Gaze x Modality | Target x Modality |
| --- | --- | --- | --- | --- | --- | --- | --- |
| ROI | **$P(BF > \log(3))$** |  |  |  |  |  |  |
| V1 | 0.43 | **1.00\*** | **1.00\*** | **1.00\*** | 0.26 | **1.00\*** | 0.92 |
| SPL5 | 0.62 | **1.00\*** | **1.00\*** | **1.00\*** | 0.29 | **0.99\*** | **1.00\*** |
| SPL7 | **0.99\*** | **1.00\*** | **1.00\*** | **1.00\*** | 0.07 | **1.00\*** | **0.99\*** |
| BA4a | **0.99\*** | **1.00\*** | **1.00\*** | **1.00\*** | 0.23 | **0.99\*** | **1.00\*** |
| PMd | 0.88 | **1.00\*** | **1.00\*** | **1.00\*** | 0.42 | **0.99\*** | **0.99\*** |
| Heschl | 0.79 | **1.00\*** | **0.99\*** | **1.00\*** | 0.03 | **1.00\*** | **0.99\*** |

When examining hand*cue modality interactions, we found no evidence of meaningful interactions in any region, indicating that representations of initial hand position were not modulated by initiation cue modality [P(log BF > log(3)) ≤ 0.42 for all regions]. A lack of differential modulation by visual or audiovisual initiation cues suggests that initial hand position representations that are less goal-relevant than target and gaze representations can be supported by the proprioceptive system, relying less on visual and/or auditory systems.

### Distinct Representations for Initiation Cue Modality

All ROIs showed distinct spatial pattern activity between trials with visual versus audiovisual go-cues [P(log BF > log(3)) = 1.00 for all ROIs]. The sensitivity to changes in initiation cue modality that we observed in all ROIs indicates that the reaches with visual go cues were represented differently than reaches with audiovisual go cues. These results support our hypothesis that sensorimotor regions would be sensitive to changes in initiation-relevant task features (Figure 3).

### Representations for Hand Position, Gaze Direction, and Target Location

The only ROIs sensitive to changes in initial hand position were BA4a [P(log BF > log(3)) = 0.99] and SPL7 [P(log BF > log(3)) = 0.99]. All ROIs were sensitive to changes in gaze direction [P(log BF > log(3)) = 1.00 for all regions] and reach target direction [P(log BF > log(3)) ≥ 0.98 for all regions], indicating distinct representations in these ROIs for left and right gaze, and left and right targets, respectively. Critically, while these contrasts do not directly test our hypotheses, including these components allows evaluating contributions of components that test our hypotheses independent of gaze, hand position, and target direction effects that are known to drive activity pattern differences in these regions^2,3,28,29^.

### Differences in Reaction Time with Unimodal and Multimodal Go-Cues

Subject showed very similar reaction times between the two testing sessions. In the audiovisual session we observed a mean reaction time of 0.762 (SD = 0.195) seconds. In the visual-only session we observed a mean reaction time of 0.765 (SD = 0.153) seconds. Paired t-tests revealed no difference between reaction time in visual versus audiovisual conditions (p = 0.947). Unlike previous literature, we did not observe a change in reaction time between conditions with visual and audiovisual go cues^30,31^. The divergence with existing literature may stem from not explicitly instructing participants to move as quickly as possible once they received a go cue. Further, many reaction time studies use simple button press tasks, whereas we used a reach task with visual cues occurring in the visual periphery. Our results suggest that changes to spatial patterned activity for target and gaze direction with audiovisual and visual cues, respectively, are not driven by behavioral differences between trials with visual initiation cues versus those with audiovisual initiation cues.

## 4. Discussion

This study investigated the effect of sensory context in which initiation-relevant cues are presented on neural representations of goal-directed reaching. Participants performed goal-directed reaches to visual targets across two sessions where reach initiation was cued with 1) only visual information, and 2) audiovisual information. We expected that sensorimotor ROIs would exhibit distinct representations for goal-relevant task features that would be modulated by initiation-relevant sensory information (i.e., initiation cue modality). Bayesian vRSAs on contrasted fMRI data provided strong support for our hypotheses, showing that goal-relevant representations in sensorimotor ROIs were largely modulated by changes in initiation cue modality.

While our main effects showing pattern dissimilarity with changes to gaze direction and target location were in line with previous literature^2,3,9,28,29,32-40^, the most notable result in the present study was that sensory context of initiation-relevant information modulates goal-relevant task representations. Specifically, the goal-relevant neural representations of gaze direction and target direction were modulated depending on whether the reaching task was cued visually or audiovisually. There are several explanations for why these sensorimotor ROIs, which are traditionally thought to encode where and how to move, would be sensitive to changes in initiation-relevant information that do not contribute to understanding the spatial structure of the task. First, several studies have indicated that regions PPC, PMd, V1, and Heschl possess neurons that respond to target stimuli across several sensory modalities^41-44^. If these regions are monitoring where/how to move as well as when to move, then any sensory change to either goal-relevant or initiation-relevant information can lead to a distinct pattern of activity.

The interaction between goal-relevant and initiation-relevant task features could also indicate that the brain is flexibly delegating these subtasks, particularly in the regions further along either processing stream. During the visual-only condition, goal-relevant and initiation-relevant information must be processed and relayed from overlapping visual areas to visually responsive parietal and premotor neurons, potentially stressing the capabilities of the visual processing stream. During the audiovisual condition, goal-relevant information is relayed via the dorsal visual stream, but initiation-relevant information can travel via the dorsal auditory stream to auditory responsive neurons downstream or the dorsal visual stream. In such a circumstance, the brain can limit the processing demand on the dorsal visual stream by allowing the dorsal auditory stream to monitor initiation-relevant sensory inputs, while the visual stream monitors goal-relevant information. The presence of a distinct representation and absence of a behavioral difference indicates the presence of neural redundancy for both goal-relevant and initiation-relevant aspects of reaching depending on sensory context.

The lack of a hand position by cue modality interaction may also be related to the use of visual and audiovisual cues to prompt movement, rather than a cue with proprioceptive stimuli. Individuals can monitor the position/orientation of their effector with visual and proprioceptive sensory information^45,46^. In the present study, the shift from visual to audiovisual did not introduce or alleviate sensory noise from proprioception. That V1 and Heschl areas showed no evidence for representations of hand position, or hand*modality interactions suggests that initial hand position representations might rely less on the visual and auditory systems. If go cues had been delivered with a combination of proprioceptive and visual information, thereby creating a conflict between tracking the go cue and tracking the effector position, we may have observed an interaction. An alternative, though not mutually exclusive, explanation for a lack of a cue modality*hand position effect is that the initial hand position is less goal-relevant than a visual target or gaze direction. As a result, the interaction with initiation cue modality is more salient for representational task features (i.e. target and gaze) that are more directly tied to the action goal. Future work should explore how different varieties of initiation-relevant stimuli interact with goal-relevant representations of movement.

Sensorimotor ROIs are usually sensitive to changes in hand position^32^; however, in the present study only regions BA4a and SPL7 showed representations of left and right starting hand position, and no regions showed a hand position by initiation cue modality interaction. The lack of support for representational dissimilarity with changes to hand position may be due to the timing in which we performed the GLM^47-52^ (i.e., closer to movement initiation). Reach direction and amplitude are processed at different periods throughout motor preparation, with reach direction being represented earlier in motor preparation and reach amplitude having stronger representations closer to initiation^47-52^. Unlike reach direction, which is directly goal-relevant, reach amplitude is a scalar value that is not directly tied to the target location. As such, representations of hand position just before movement initiation may be less goal-relevant than representations of gaze or target direction.

Understanding the complex interactions of sensory and motor pathways in the production of goal-driven movement may provide benefits for individuals with neurologic disorders or insults. For instance, individuals with Parkinson’s Disease (PD) show altered movement initiation during both reaching^53^ and locomotor tasks^54,55^, and poor ability to use visual information to plan precision stepping tasks (i.e., a goal-directed reach task with the lower limbs)^56,57^. Experiments using external cues have shown improvements in movement execution with external sensory cues^55,58-62^and improvements in movement preparation with external auditory cues^54,58^. To our knowledge, there has been limited study of the effects of multimodal cues on feedforward gait behavior in persons with PD. While speculative, supplying a multimodal sensory cue at gait initiation may provide a level of redundancy that reduces the burden on the visual system and improves behavior. Future work should explore how representational similarity with different sensory cues changes in populations with neuromotor deficits, such as persons with PD.

One potential limitation of this study relates to the shorter distance between the two starting hand positions than the two target and two gaze positions. Parietal and motor areas are sensitive to changes in gaze, target, and effector distances during reaching tasks ^45,49,63^. The left- and right-hand positions were separated by 2.3 cm, whereas target and gaze locations were separated by 6.9 cm and 16.1cm, respectively. This small distance between left- and right-hand positions may partly explain why we observed few distinct neural representations for the effect of starting hand position and no interaction between hand position and cue modality. However, since other studies have seen effects with similar distance reaches^9,37,49^, it is likely that the direction of initial hand position is less represented than goal-relevant target and gaze positions in the lead-up to movement initiation. The narrow orientation of the hand and target buttons was selected to limit movement and allow participants to reach from the hand to the target buttons without requiring movement of the entire arm. We also chose target and gaze positions such that targets on all trials would always fall within an observer’s near peripheral zone within which there are little changes in in visual acuity^64^. In this way, we minimized between-condition activation pattern differences related to visual acuity differences.

Another potential limitation also stemming from a design choice might have influenced a lack of a behavioral difference between audiovisual and visual conditions, unlike that shown in previous multisensory facilitation studies. In those studies, the audiovisual conditions are the sum of stimuli from distinct auditory and visual conditions^30,31^. Our audiovisual condition featured two initiation cues: a visual initiation cue (i.e., the target light) and an audio-initiation cue. In our design, we opted to match the number of initiation cues between audiovisual and visual conditions, such that any activation pattern differences were driven only by a difference in the initiation cue modality rather than the number of cues given to initiate the movement. Thus, like the audiovisual condition, the visual condition featured two visual initiation cues (i.e., the gaze light and the target light). Adding an auditory cue to the visual condition would have created an imbalance between the total number of initiation cues in our contrasted conditions (i.e., two for the visual condition and three for audiovisual). We suspect that such a methodological choice likely would have elicited reaction time differences due to multisensory facilitation. However, it would also have complicated the interpretation of patterned neural activity as any differences could have been attributed to the change in the quantity of initiation cues or the modality of initiation stimuli. The present design allowed us to directly test how pattern differences change in the presence of initiation cues from visual versus auditory and visual channels. Finally, the distinct representations we observed may also be explained by the gaze light changing color in the visual condition but not the audiovisual condition. Visual areas are sensitive to both auditory stimuli and color stimuli ^65-67^. However, the pathway responsible for processing color passes from V1 through the inferior temporal cortex ^65,68,69^. Sensorimotor and superior temporal regions, where we also observed these interactions, are more commonly associated with sensitivity to polysensory stimuli^41,43^ than simple color changes ^65,67^.

## 5. Conclusion

Altogether, our results show that representations of the task features of goal-directed reaching differ depending on the sensory context of initiation-relevant information.

Representations of visual task features, such as gaze direction and target location, were particularly sensitive to changes between visual and audiovisual cueing. Future work should examine the effects of other multimodal go-cues, such as combined proprio-visual or proprio-auditory, on the task feature representations of goal-directed reaching, and also how different groups with neuromotor deficits incorporate initiation-relevant information for goal-directed tasks.

## 6. Acknowledgments

The authors would like to thank Ale Harris Caceres and Karla Barajas for their assistance with data collection.

